# An olfactory pattern generator for on-demand combinatorial control of receptor activities

**DOI:** 10.1101/2021.05.31.446433

**Authors:** Guangwei Si, Jacob Baron, Yu Feng, Aravinthan D.T. Samuel

## Abstract

Olfactory systems employ combinatorial receptor codes for odors. Systematically generating stimuli that address the combinatorial possibilities of an olfactory code poses unique challenges. Here, we present a stimulus method to probe the combinatorial code, demonstrated using the *Drosophila* larva. This method leverages a set of primary odorants, each of which targets the activity of one olfactory receptor neuron (ORN) type at an optimal concentration. Our setup uses microfluidics to mix any combination of primary odorants on demand to activate any desired combination of ORNs. We use this olfactory pattern generator to demonstrate a spatially distributed olfactory representation in the dendrites of a single interneuron in the antennal lobe, the first olfactory neuropil of the larva. In the larval mushroom body, the next processing layer, we characterize diverse receptive fields of a population of Kenyon cells. The precision and flexibility of the olfactory pattern generator will facilitate systematic studies of processing and transformation of the olfactory code.

## Introduction

Sensory systems like vision and hearing encode complex stimuli in the activity of sensory neuron arrays. Progress in understanding these systems has benefited from stimulus methods that span the coding capacity of sensor arrays. However, the number of different inputs that might be coded grows rapidly with array size. With visual images or sounds, it is still straightforward to generate and deliver complex stimuli – e.g., combinations of primary colors, pixels or sound frequencies – with independent control over all individual sensory neuron types. In olfaction, each odorant stimulus is associated with a specific combination of activated olfactory receptor neurons^1^. However, olfactory receptors are not tuned by any continuous physical properties of odorants in the same way as color in vision or sound frequency in hearing. Thus, it has been difficult to deliberately compose odor stimuli to precisely and flexibly target any desired combination of receptors.

An ideal method for olfactory stimulation would allow the experimenter to activate any ORN combination on demand. One way to span the combinatorial possibilities of a sensory code is to project combinations of “primary inputs” onto a sensor array, analogous to the way that any hue can be generated with three primary colors. An ideal primary input activates only one sensory neuron type without cross-talk. A mixture of primary odorants for a set of ORNs would allow the experimenter to specifically activate all ORNs in that set.

Here, we develop an olfactory pattern generator that enables us to probe the combinatorial possibilities of the olfactory code in the *Drosophila* larva. The larva has a compact olfactory system with a small number of olfactory receptors^2,3^, an ideal system to develop and demonstrate the principles of an olfactory pattern generator. We identify a set of primary odorants for the *Drosophila* larva and pinpoint the optimal concentration for most of its ORN types. We controllably deliver these primary odorants using microfluidics, which is particularly effective for studying the chemosensory system of small intact animal like *C. elegans*^4^ and *Drosophila* larva^5^. Our microfluidic device is capable of mixing and delivering any set of primary odorants at their optimal concentrations (Fig.1A).

**Figure 1.**
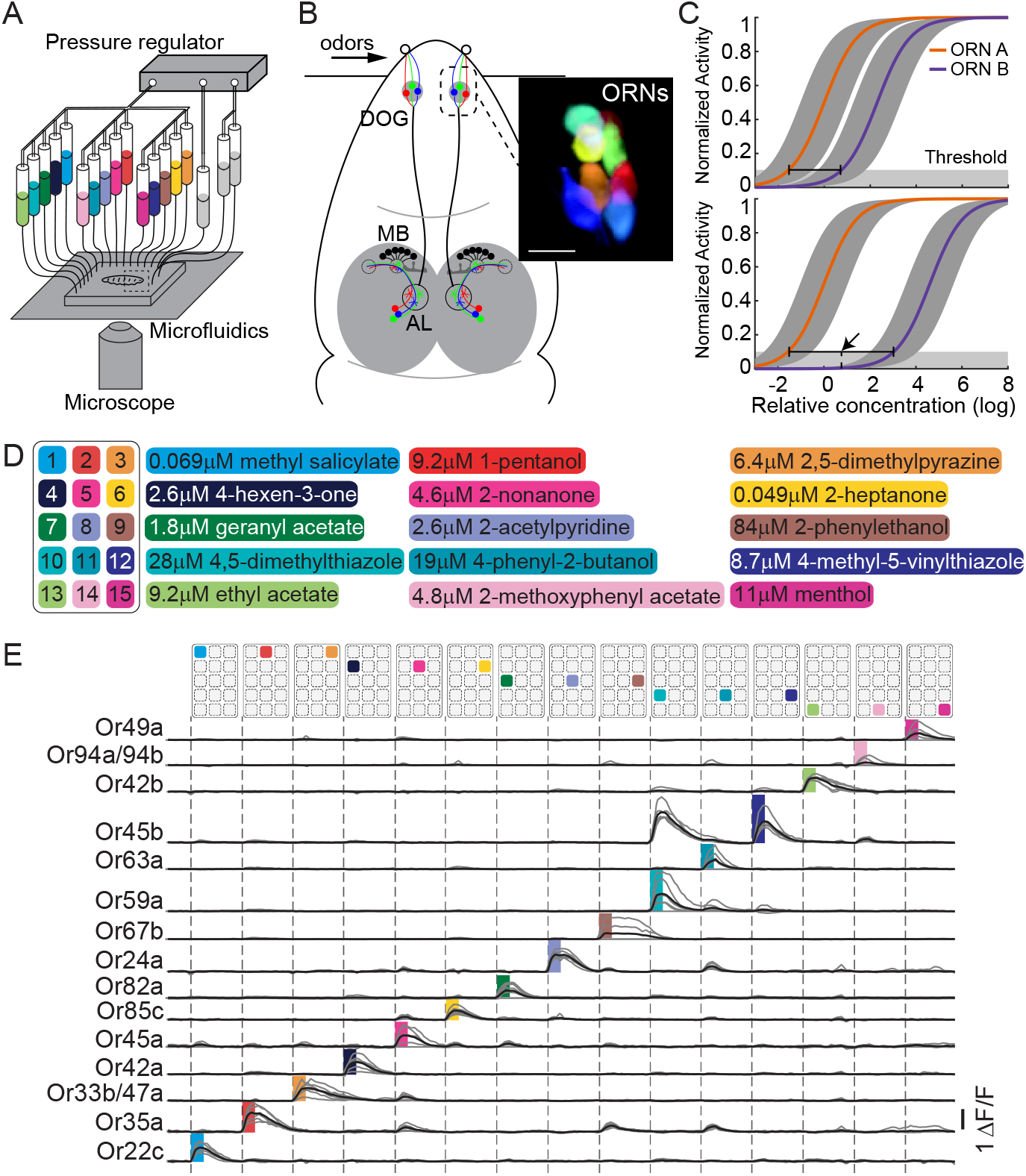
Experiment setup and selection of primary odorants. A. Schematic of the experiment setup. A pressure regulator outputs three independent adjustable pressure sources and drives odorant solution flow through a microfluidic chip. Odor-evoked neural activities from an immobilized first instar larva are recorded using a confocal microscope. B. Schematic illustration of the larval olfactory circuit. ORNs, AL, and MB form the first three layers of olfactory circuit. Inset, ORNs segmented based on their activity patterns (indicated by colors). Scale bar, 10 *μm*. DOG, dorsal organ ganglion; AL, antennal lobe; MB, mushroom body; ORNs, olfactory receptor neurons. C. Comparing candidate primary odorants (top and bottom) and identifying the optimal concentration. Consider the dose-response curves of the two most sensitive ORNs (ORN A and ORN B). The further these two curves apart, the better the separation in their activities (bottom panel). The gray regions indicate the variance of the sensitivities between the odorant and ORN. The optimal concentration lies at the midpoint (arrow head) between these two curves crossed by the activation threshold (TH.). D. List of the name and the optimal concentration of selected primary odorants. E. Calcium transients in 15 ORN cell bodies in response to pulsed presentations of the primary odorants.

In the larva, the olfactory pattern generator can be used with optical imaging to measure olfactory representations and processing throughout the circuit. We demonstrate its effectiveness by examining how different combinations of activated ORNs are mapped to two new forms of olfactory representation at successive circuit layers^6^. We study a prominent interneuron in the antennal lobe and populations of Kenyon Cells (KCs) in the mushroom body (Fig.1B). The olfactory pattern generator allows us to study how olfactory representations are transformed across layers of an olfactory circuit.

## Results

### Primary odorants at optimal concentrations target individual ORNs

The first step in building an olfactory pattern generator is to collect a set of primary odorants. Each primary odorant should reliably activate only one type of ORN. However, the number of activated olfactory receptors varies with odor concentrations, e.g. odors at higher concentration tend to activate more ORNs^7,8^. Any primary odorant will only work within a specific concentration range. The odorant concentration should be high enough to activate the most sensitive ORN, but not too high to activate the second most sensitive. Consider the dose-response relationships of a candidate primary odorant for the two ORNs that are most sensitive to it (Fig 1C). The farther apart the two dose-response curves are from each other, the better that candidate odorant might work as a primary odorant. The optimum concentration that maximizes the chance that only the most sensitive ORN is activated above a threshold lies at the midpoint between where the two dose-response curves cross that threshold (see Methods, Fig. 1C).

We applied our strategy to define primary odorants using a recently acquired larval odor-ORN response dataset^5^. We were able to identify a set of 14 primary odorants targeting 14 different ORNs and their optimum concentrations (see Methods, Fig. 1D). We included one odorant in our panel, 4-methyl-5-vinylthiazole, that reliably activates two ORNs with comparable magnitude at its optimum concentration. To verify the properties of these primary odorants, we expressed the calcium indicator GCaMP6m^9^ in all larval ORNs under the control of *Orco-Gal4*^10^, and imaged ORN calcium dynamics in response to pulsed odor stimuli. We verified that all primary odorants individually activate their target ORNs with minimal cross-talk to other ORNS (Fig. 1E).

### On-demand mixing primary odorants generates ORN activity patterns

The next step is to mix any combination of primary odorants at their optimal concentrations to stimulate any desired pattern of ORN activity. Mixing solutions leads to dilution – equally mixing *N* odorants dilutes their original concentrations to 1/*N*. Our system needs to compensate for the effect of dilution when mixing different number of primary odorants. We implemented our mixing setup with dilution compensation using microfluidics (Fig. 2).

**Figure 2.**
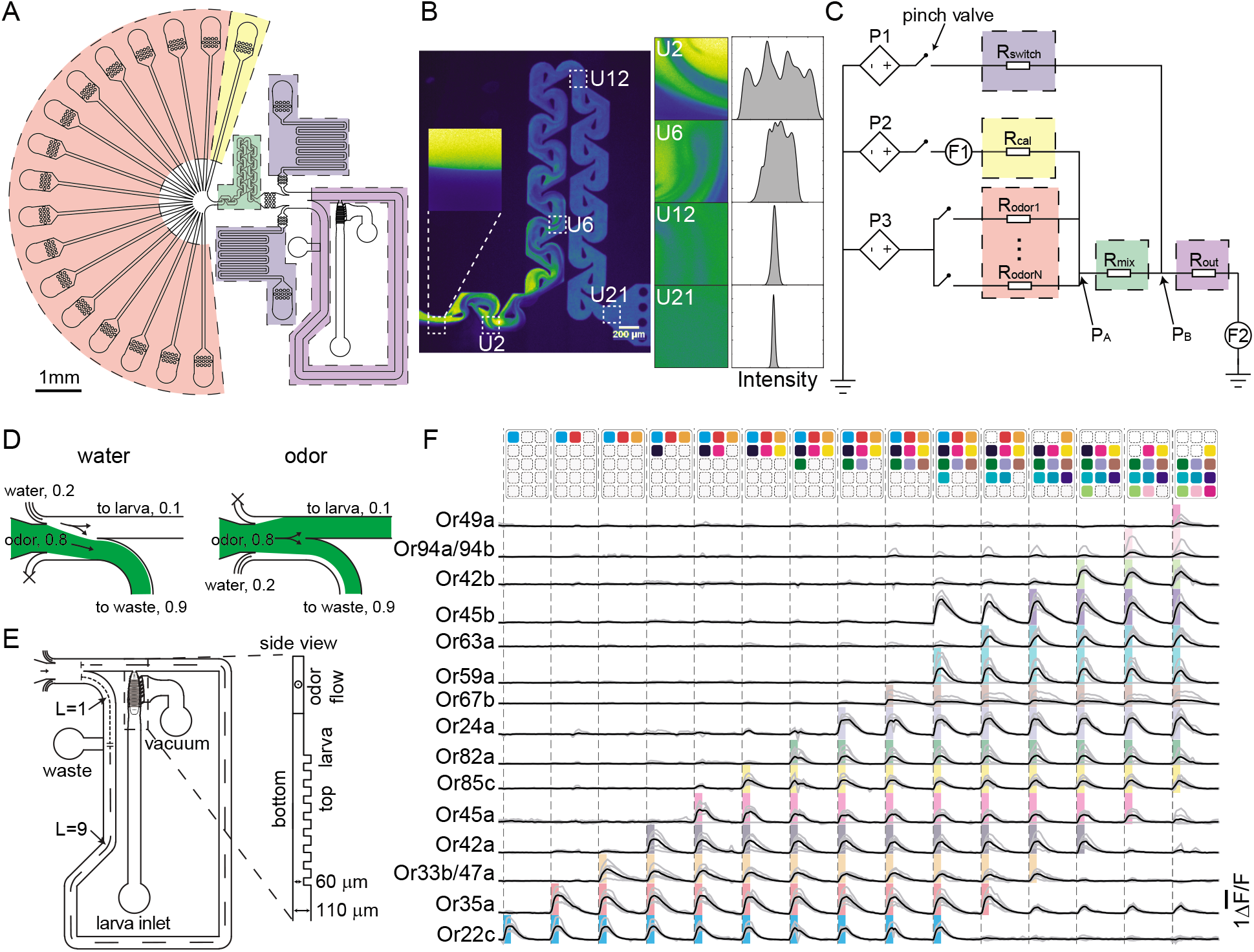
Mix primary odorants to generate ORN activation patterns using microfluidics. A. Layout of the microfluidic chip. The orange, yellow, green, gray, and purple regions indicate odorant input channels, dilution channel, mixer, switch channels and outlet channel, respectively. B. Visualizing the mixing process through the Tesla valve structures using fluorescent dye. After 20 units, the two streams are well mixed. C. Schematic diagram of the control circuit to regulate the flow rates in each channel. P1-P3 are programmable pressure controllers. F1, F2 are flow meters. Color shaded regions indicate the flow resistances of the corresponding colored channels in panel A. D. Switch between water and odors. Open the top channel and close the bottom one to direct water flows to the animal. Flip the ON/OFF states of these two channels to switch between water and odorants. The numbers indicate the volume ratio occupied by each stream. E. Outlet channel and the larval channel. The side view of larval channel shows the patterned ridges used to immobilize to larval body. F. Time course of 15 ORNs’ responses to a series of mixed primary odorants.

We designed a microfluidic chip with input channels for each primary odorant that can be selected to be included in mixtures (channels shaded in orange, Fig. 2A). The flows from all open channels are collected and mixed using a passive in-plane micromixer (the green region, Fig. 2A). The micromixer consists of a series of reversed Tesla valves structure. A Tesla valve mixer produces transverse dispersion by the Coanda effect and homogenizes input fluid streams^11^. Imaging the mixing of a fluorescent and non-fluorescent stream shows that, after passing through 20 Tesla valves in series, the two streams are well mixed (Fig. 2B). We set the input concentration of each primary odorant to be *N*× its optimal concentration, then adjust the flow from a dilution channel (the yellow channel in Fig. 2A) to bring the final concentration of each odor component to 1/*N* of the input concentration. High-precision flow meters and pressure controllers are used to monitor and adjust the flow rates of the input and output channels (Fig. 2C). A PID controller keeps the output flow constant by tuning the flow rate of the dilution channel while varying the number of mixed primary odorants (see Methods). After the solution is well mixed, we control the ON/OFF of two switch channels (gray channels, Fig. 2A) to switch between water and mixed odorant (Fig. 2D).

With our panel of primary odorants and the mixer, we are able to deliver diverse stimuli to the *Drosophila* larva. To apply as many stimuli as possible to each animal in each experiment, we improved the immobilization method for long-term imaging of a live larva. We designed ridges interdigitating the larval body segments that inhibit expansion and contraction during peristalsis (side view, Fig. 2E). In addition, we used a side channel connected with negative pressure to hold the larva body by suction. These techniques produced stable and robust recordings with negligible motion for up to 40 minutes.

To demonstrate that the system works as intended, we imaged calcium dynamics in all 21 ORNs in response to mixtures of primary odorants with different numbers of components. When we deliver different combination of primary odorants, the larval ORNs show expected activation patterns. Individual ORN responses were comparable whether a primary odor was delivered by itself or with any combination of other primary odors (Fig. 2F).

#### Spatially structured response in a local interneuron

To demonstrate the utility of the method, we applied it to investigate how olfactory representations progress from the ORN layer to downstream circuits. The insect antennal lobe has a glomerular structure: each glomerulus receives input from a specific ORN and sends output via projection neurons to downstream circuits. The wiring diagram of the larval antennal lobe reveals several interneurons that innervate many or all glomeruli^12,13^. In both adult and larva, inhibitory interneurons are thought to provide broad feedback inhibition to most or all glomeruli^14–16^. However, the role of feedback inhibition in the antennal lobe may be more complex^17^. One potential source of complexity would be any spatial structure in the interneuron activity pattern that depends on olfactory inputs. We explore this possibility by imaging the spatial activity patterns of one prominent multiglomerular interneuron called Keystone in response to odors and odor mixtures^12^.

We studied Keystone by cell-specific expression of GCaMP6m using the Gal4 line GMR28F06^18^ (Fig. 3A). This enabled us to image the full spatial extent of its neuropil extending throughout the antennal lobe. When we subjected the larva to a large panel of pure odors and mixtures, we observed substantial stimulus-specific spatial heterogeneity in dendritic calcium dynamics (Fig. 3). One way to identify distinct dendritic response regions is to cluster voxels by their time-series activity patterns. We used t-SNE to group voxels based on the similarity of these time-series^19^ (Fig. 3C, see). In each animal that we studied, t-SNE revealed ~10 regions of the dendritic field with different odor responses (Fig. 3B). Some regions were roughly the size of individual glomeruli. Other regions likely spanned several glomeruli. Each region is likely to reflect the spread of calcium dynamics from activated dendritic branches into nearby neuropil. If so, neighboring regions might be more correlated by local spread of calcium dynamics than well-separated regions. To test this, we computed the pairwise similarity in the activity of different regions identified by t-SNE in response to the stimulus panel. Indeed, the similarity in the activity of two regions is positively correlated with the distance of spatial separation within the antennal lobe (Fig. 3D).

**Figure 3.**
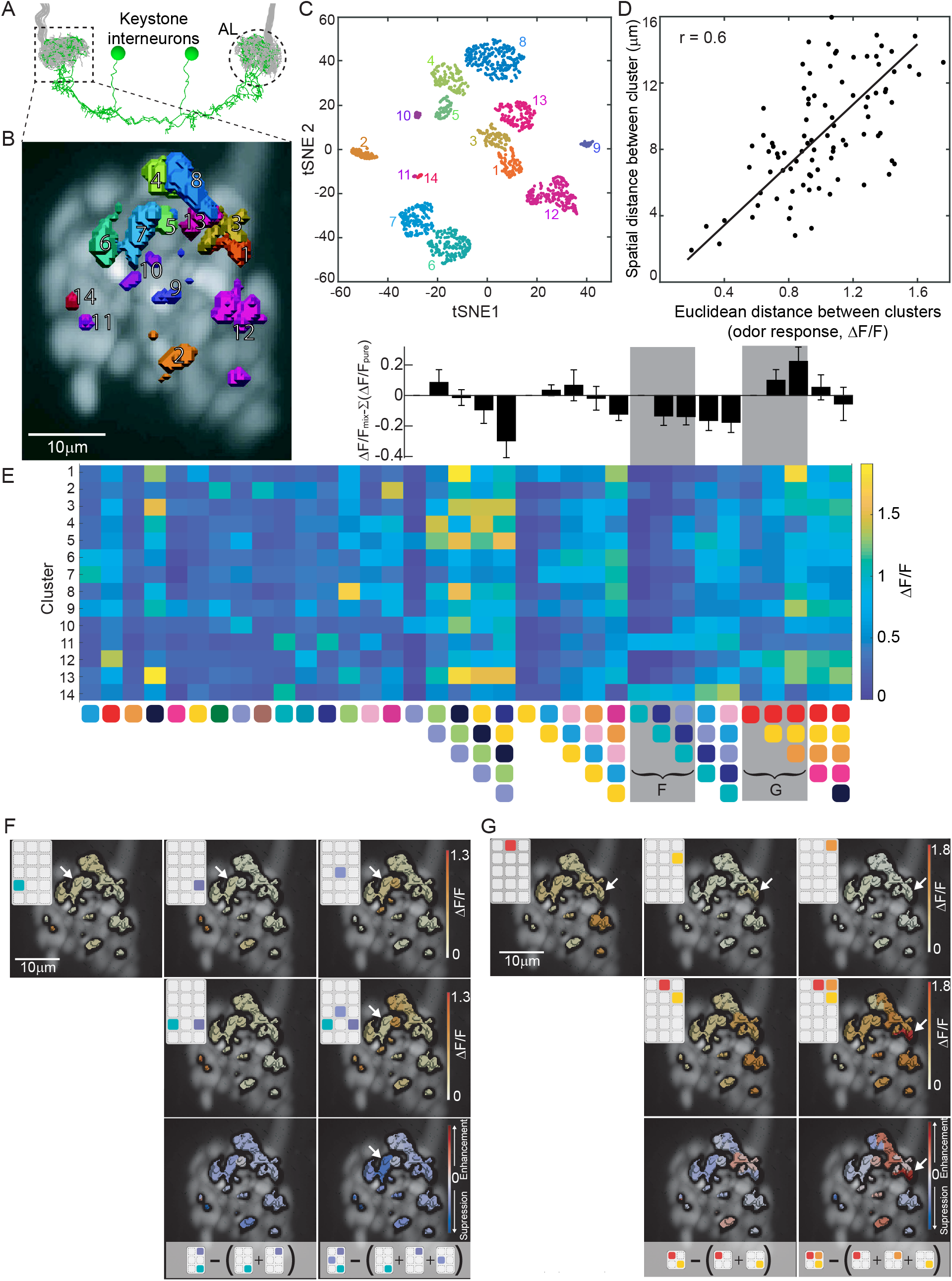
Compartmentalized calcium activity and integration of ORN inputs by an interneuron in the AL. A. Anatomy of a pair of Keystone interneurons. The neuropils innervate bilateral AL and receive inputs from ORN axons (gray). The calcium activity from Keystone neuropil within one AL. B. Compartmentalized calcium activity of the Keystone interneuron. The foreground depicts compartments of clustered activities within the Keystone interneuron’s neuropil. Number and color of the compartments correspond to clusters in the tSNE space in panel C. The background is RFP-labeled ORN neuropils, illustrating the shape of AL. C. Clustering the voxels in the tSNE space indicates the temporal correlation of voxel’s response signal. D. Correlation of cluster’s pairwise distances measured in the physical space and the activity space. E. Heatmap of the Keystone intereuron (shown in panel B) cluster’s activity responding to the primary odors and their mixtures. On top of the mixture panels are averaged comparisons between the actual response to the mixture, versus the linear sum of the response to individual primary odorants (N = 5). F. G Illustration of the different nonlinear additive effects on the Keystone interneuron, in responding to two different groups of primary odors. The first row shows the activity pattern in response to the pure primary odors. The second row shows the patterns in response to two and three mixtures. The third row shows the relative difference between the actual mixture responses and the summed pure responses. Clusters showing significant differences between these two groups are highlighted by the arrows.

We compared the spatial structure of Keystone activation to pure odors and mixtures. As odors are added to a mixture, more ORNs will provide excitatory input to regions of its dendritic tree. As we increased the number of odors in a mixture, we observed an increase in the total amplitude of integrated calcium dynamics over the dendritic field (Fig. 3E). We found that the response to mixtures is not a simple sum of the individual responses to each mixture ingredient delivered one at a time (Fig. 3D). We found that mixtures comprised of aliphatic molecules, for example, could evoke a superlinear response in some regions of the dendritic field but a sublinear response in others (Fig. 3F). In contrast, mixtures of aromatic molecules evoked sublinear responses throughout their dendritic field (Fig. 3G). We estimate superlinearity (or sublinearity) as when the response to a mixture is larger (or smaller) than the summed responses to each ingredient delivered one at a time.

The spatial structure of calcium dynamics in the dendritic field of the Keystone interneuron carries information about the olfactory input. Keystone responds differently to different types of odorant molecules. Superlinear and sublinear integration of ORN inputs in the spatial structure of its activity are suggestive of different nonlinear filters in the interneuron layer of the antennal lobe. Feedback inhibition in the antennal lobe may be stimulus-dependent in ways that support more complex functions.

#### Diverse receptive fields across Kenyon Cells in the mushroom body

Next, we used our setup to measure olfactory representations in an ensemble of downstream neurons. The insect mushroom body, a center for learning and memory, is thought to recode olfactory inputs in its Kenyon Cell (KC) population^20,21^ (Fig. 4A). Each KC receives input from a subset of PNs from the antennal lobe^22–25^ and spikes only when enough PN inputs are active^26,27^. Thus, each KC may act as a feature detector for a specific set or combination of PN activities. Understanding the distribution of tuning properties among KCs is essential to understanding the olfactory code of the mushroom body.

**Figure 4.**
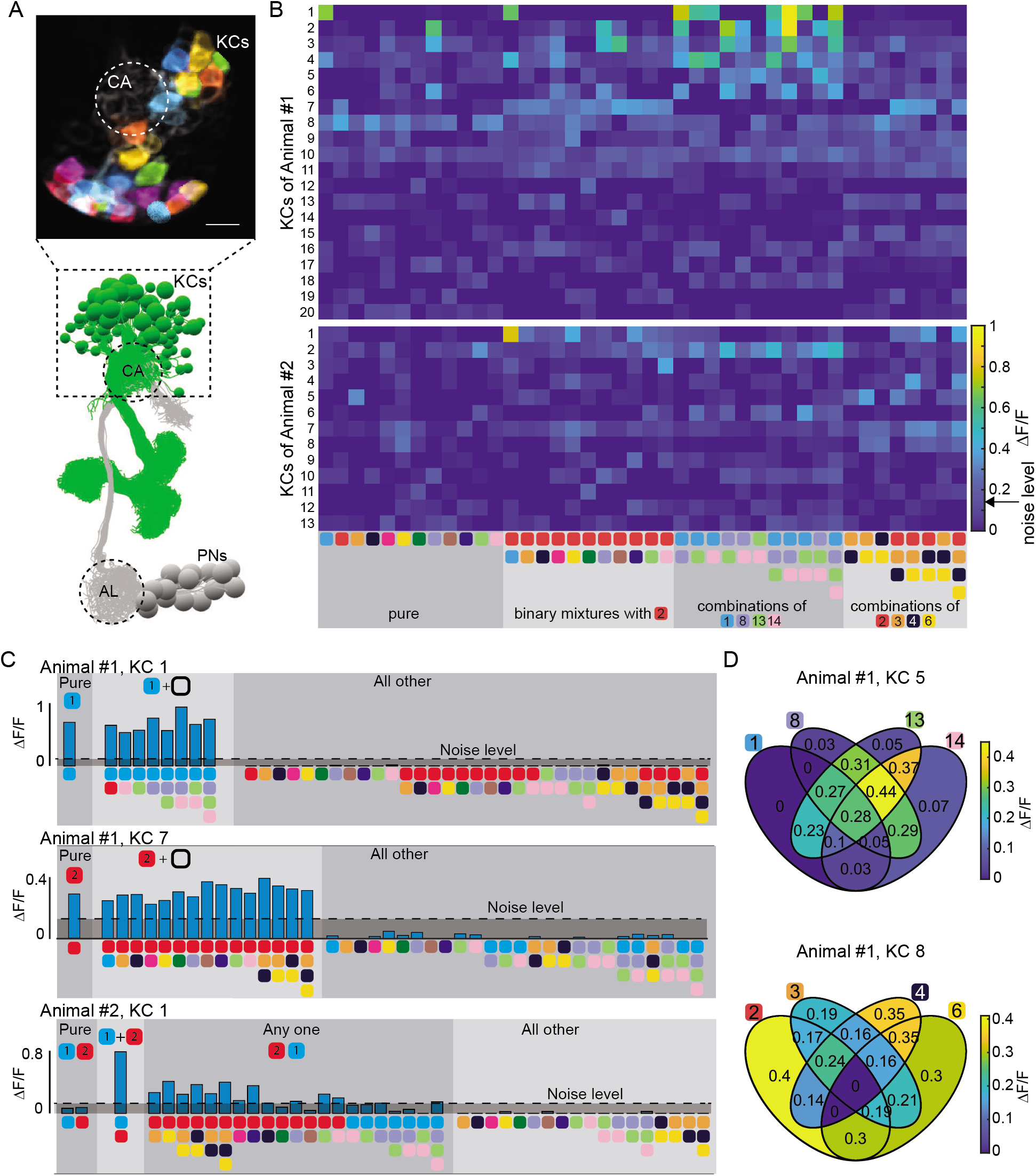
Diverse responses profiles across a population of Kenyon Cells in the mushroom body. A. Illustration of the circuit anatomy of the Kenyon cells in the mushroom body. KCs, Kenyon cells; CA, Calyx; AL, antennal lobe; PNs, projections neurons. Calcium imaging was performed on the soma of the Kenyon cells. B. Heatmap of KCs’ responses to four groups of odorants, pure, binary, and two groups of complete combinations of four primary odorants. The data were from two animals, KCs were selected for at least one response above the noise level, which is indicated on the color bar. C. Example KCs activated by selective combinations of the primary odors. i and ii show two KCs response to a certain pure primary odorant and mixtures containing that odorants. iii, one example KC shows strong response only to a binary mixture, weak or negligible responses to individual components and other odorants. D. Projection of two example KCs’s activity onto a Venn-diagram, showing center surrounded receptive field. The top one resembles an ON center neuron and the bottom resembles an OFF center neuron.

Using the olfactory pattern generator, we designed two types of stimulus panels to characterize KC response properties. One panel was designed to broadly sample the olfactory receptor neurons – we individually delivered every primary odor in our library as well as every 2-odor combination containing 1-pentanol and another primary odor. Another panel was designed to sample all possible combinations within sets of four ORNs – we delivered every 1-, 2-, 3-, and 4-odorant combination (15 in total) of the primary odorants associated with the 4 ORNs. For one subpanel, we chose 4 ORNs that are more sensitive to aliphatic odorants For the other, the chosen ORNs are more sensitive to aromatic odorants (Fig. 4B).

We recorded somatic calcium dynamics in the ~100 visible KCs of first instar larvae expressing GCaMP6m under the control of 201Y-Gal4^28^. In each animal that we studied, we found that ~20 KCs displayed a calcium response to at least one stimulus from one of the panels (Fig. 4B). These KCs showed a remarkable diversity of response types. We found a subset of KC with bimodal responses, strongly responding to some stimuli but negligibly responding to others. The simplest “bimodal KCs” responded to one of the primary odors delivered individually and equally well to any mixture that contained that primary odor. For example, the KC shown in Fig 4Ci responds equally well to methyl salicylate or any mixture containing methyl salicylate. Such KCs might represent the single-claw KCs discovered in the connectome that receive input from exactly one PN^25^. These KCs resemble relays that pass a signal from single presynaptic PNs to their postsynaptic partners.

Other KCs responded only to mixtures containing multiple primary odors. Some KCs were activated by a minimum set of primary odors. For example, the neuron shown in 4 Ciii is active whenever methyl salicylate and 1-pentanol are presented together but not individually. Such KCs might represent the two-claw KCs discovered in the connectome that receive inputs from two PNs^25^. These KCs resemble “AND” gates in requiring two simultaneous input signals - only when both inputs are on will the output be on^26,27^.

Many KCs exhibited response properties that were more complex than relays or simple logic gates. Some did not have ON-OFF bimodal responses, but instead had graded responses to different inputs. With our 4-odor panels, we measured the response in each KC to all 15 different stimuli that can be made from 4 primary odors. Every stimulus that can be made from four primary odors can be represented as a region in a Venn diagram. Illustrating the response evoked by each input with a Venn diagram reveals the particularly complex response properties of some KCs (see Fig.4D, S1 and S2). For example, the KC shown in Fig 4D upper panel, exhibits negligible or weak responses to any one primary odor, but stronger responses to mixtures with multiple odors. Responses to mixtures were also variable, with some 2-odor mixtures evoking stronger responses than others. The total KC response is not a simple sum of excitatory inputs, as this KC responds more effectively when subjected to some 2- and 3-odor mixtures than to the 4-odor mixture.

Surprisingly, some KCs could be inhibited when more primary odors were added to a mixture. For example, the KC depicted in Fig. 4D lower panel responded most strongly to single primary odors, and less for mixtures with more odors. For this KC, the weakest response was evoked by the 4-odor mixture. This KC appears to have a low response threshold to individual odors, but is significantly inhibited by the addition of other odors.

Depending on the overall composition of a mixture, each primary odor can be viewed as having either an inhibitory or excitatory effect on a KC, and these effects can vary in strength. Varying levels of KC excitation may be due to differences in the strength of direct presynaptic inputs from the antennal lobe or differences in the activation thresholds of each KC. The connectome suggests that odor-dependent inhibition may be due to negative feedback within the mushroom body. The main negative feedback is from the anterior paired lateral (APL) neuron that receives excitatory input from all KCs and provides synaptic lateral inhibition to all KCs^29^.

## Discussion

We have developed and validated a method to generate odor stimuli that is well suited to analyze combinatorial codes in olfaction. We implement this method using microfluidics to study the olfactory system of the *Drosophila* larva. Our method allows an experimenter to mix and deliver any set of primary odors to the *Drosophila* larva on demand, evoking the activity of any desired combination of ORNs.

Optogenetics has recently been used for combinatorial control of olfactory inputs using patterned illumination^30–32^. This is done by shaping each light pattern for stimulation to the physical layout of an olfactory circuit, such as the glomeruli of the insect antennal lobe or mouse olfactory bulb. However, each light pattern typically has to be programmed in each experiment to accommodate anatomical differences between animals.

Mixtures of private odors – odorants that specifically activate single olfactory receptor types with minimal cross-talk – are a natural and flexible way to control olfactory inputs. The small size of the *Drosophila* larva olfactory system has allowed the identification of private odors for most of its ORNs^33^. Our method uses a set of private odors as primary odorants, analogous to the way that three primary colors can be used to create all hues seen by a trichromatic visual system. Furthermore, we use this set of primary odorants to study olfactory representations to combinatorial patterns of ORN activation. The olfactory pattern generator is able to project any desired activation pattern onto the ORN ensemble to probe their olfactory representations.

We discovered a new type of complexity in the response of a broad local interneuron in the antennal lobe. The antennal lobe is the first olfactory relay in *Drosophila*, formatting olfactory information to be used by other brain regions. For example, broad local interneurons within the antennal lobe provide negative feedback inhibition that is useful for normalization, reducing the dependence of antennal lobe output on total odor concentration. Here, we studied a broad local interneuron called Keystone that was identified in the wiring diagram of the *Drosophila* larva’s antennal lobe^12^. When we stimulated the larva with a broad set of odors and odor mixtures, we discovered spatial structure in the odor-evoked calcium dynamics of Keystone’s dendritic field. The olfactory representation encoded in the Keystone dendritic calcium dynamics is more complex than a sum of olfactory inputs, and may contribute to odor-dependent feedback inhibition and filtering of sensory information.

We also examined ensemble-wide representations in the Kenyon Cells (KCs) of the mushroom body, a learning and memory center. KCs greatly outnumber ORNs, and a typical olfactory input activates a much smaller fraction of KCs than ORNs. Sparse coding by KCs is thought to result from their random integration of small numbers of projection neuron (PN) inputs from the antennal lobe. Each PN that innervates the mushroom body is activated by one ORN type. We found that the ensemble-level activity of the KC population is not only sparse, but that individual KCs can have surprisingly complex properties. KCs are not simply activated by minimal sets of odorants, but can have complex receptive fields that suggest excitation by some odors and inhibition by others. Such responses are reminiscent of cells with classical center-surrounded receptive fields in the early visual system that exhibit ON and OFF tuning to different parts of input space^34^. Feedback inhibition might only sparsify the KC code, as has been previously suggested, but might contribute to the complex receptive fields of individual KCs. The major source of feedback inhibition in the mushroom body is the APL interneuron that receives input from all KCs and provides negative feedback to all KCs. A richer diversity in KC response types, mediated by complex patterns of feedback inhibition, might increase the sophistication of the population-level code of the larval mushroom body.

We have shown that an olfactory pattern generator is useful for discovering and analyzing new complexities in the olfactory representations of combinatorial patterns of ORN activity. The use of an olfactory pattern generator may be extended to other animals. The first step is to build a catalogue of private odorants for each ORN that can be leveraged as primary odorants for mixtures. Once primary odorants have been determined, they must be implemented in a mixture and delivery system that maintains the optimal stimulus concentration of each odorant while minimizing cross-talk. Microfluidic systems are appropriate for aqueous delivery of odorants to animals like *C. elegans*, *Drosophila* larvae, and zebrafish. For animals like adult flies or vertebrates, the recent development of gas-based olfactometers with high flexibility^35^ and dynamic control^36^ allows the construction of similar pattern generators. Olfactory pattern generation with primary odorants is an ideal way to probe combinatorial codes in olfaction.

## Methods

### Drosophila stocks

The following transgenic stocks were obtained from the Bloomington Drosophila Stock Center(BDSC) and used in this study: UAS-mCherry.NLS; UAS-GCaMP6m (*UAS-mCherry.NLS*, BDSC#38425; *UAS-GCaMP6m*, BDSC#42750), UAS-GCaMP6m; Orco::RFP (*UAS-GCaMP6m*, BDSC#42748; *Orco::RFP*, BDSC #63045), UAS-mCD8::GFP; Orco::RFP (BDSC#63045), Orco-Gal4 (BDSC#23292), GMR28F06-Gal4(BDSC#48083), 201Y-Gal4(BDSC#4440). All calcium imaging experiments were performed on first instar larvae of either sex.

### Identifying primary odorants and optimal concentrations

In larval *Drosophila*, ORN odor’s dose-response relationship could be described by a Hill function, 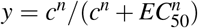, here the amplitude is normalized to 1, *c* is the odor concentration and *EC*_50_ is the half maximal effective concentration^5^. For convenience, we use the logarithm of odor concentration, *x* = *ln*(*c*), *k* = *ln*(*EC*_50_), *y* = 1/(1 + *e*^−*n*(*x*−*k*)^). To evaluate a candidate primary odorant and determine the optimal concentration, we consider the two most sensitive ORN’s responses, *y*_1_ = *y*_1_(*k*_1_) and *y*_2_ = *y*_2_(*k*_2_) (‘ORN A’ and ‘ORN B’ in Fig. 1C). The criteria of strongly activating the most sensitive ORN and not activating the second most sensitive one is quantified as *y*_1_ − *y*_2_. This term reaches a maximal value at *x* = (*k*_1_ + *k*_2_)/2, and the value increases monotonically with Δ*k* ≡ *k*_2_ − *k*_1_. The further apart these two dose-response curves, the better the candidate will work as a primary odorant. The optimal concentration is located at the midpoint between the two curves. The conclusion holds when we use other objective functions, e.g., *y*_1_ * (1 − *y*_2_). Furthermore, when we consider the variation of the sensitivities, e.g., if *k* follows a Gaussian distribution, the conclusion also holds and the objective function reaches its maximum value with the highest likelihood.

When mixing multiple primary odorants, the non-primary activation (i.e. the weak responses of the second most sensitive ORNs) may accumulate and generate significant unexpected responses. To avoid this, we tended to adopt lower activation thresholds. We empirically chose 10% of the ORN maximum response as the activation threshold when using mixtures. In this case, the optimal concentration (in logarithm scale) could be modified as *x* = (*k*_1_ + *k*_2_)/2 + *ln*(9)/*n* (arrow in Fig.1C bottom panel).

Our recent study^5^ mapped the dose-response relationship between 34 odorants and all the 21 larval ORNs. We rank the 34 odorants by the difference of sensitivities to the two most sensitive ORNs, i.e. Δ*k*. The larger the value of Δ*k*, the better a candidate primary odorant. When Δ*k* is small enough, 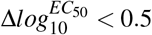, we reject that odorant as a candidate primary odorant. For a given ORN, if there are multiple candidates, i.e. that ORN is multiple odorant’s most sensitive ORN, we choose the odorant having the largest Δ*k*.

### Microfluidic chip design, fabrication and calibration

AutoCAD (Autodesk, Inc.) was used to design the microfluidic chip shown in Fig. 2B. The layout consists of two layers: the first layer includes the majority of the features and was fabricated with a height of 60*μm*, which is optimized for maximizing the mixing efficiency of the Tesla valve mixer. The second layer mainly includes the larva channel for immobilization (Fig. 2E), with a height of 100*μm*, which fits the body thickness of the first instar larvae.

Fabrications of the master and polydimethylsiloxane (PDMS)-based microfluidic chips follow standard photolithography and soft lithography protocols. The reuse of the microfluidic chip was restricted across odorant sets and across days to prevent cross-contamination of the odorants.

Fluorescnt dye was used to calibrate the mixing process of the Tesla units. The flow ratio of water and fluorescent was set as 1:1, and the total flow was set in the range of the operating flow rate. After each Tesla unit, the distribution of the pixel-wise fluorescent intensity was measured to quantify the mixing process. The narrower of the intensity distribution indicates a more uniform mixture.

### Regulate the dilution

Precise odorant dilutions are achieved using three high precision pressure controllers (MFCS-EZ, Fluigent, Inc.) and two flow meters (Flow Unit, Fluigent, Inc.) as shown in Fig. 2C. Pressure controller *P*1, *P*2, and *P*3 regulate the flow of primary odorant inputs, dilution channel, and the switching channels, respectively. Flow meter *F*1 and *F*2 measure the flow rate of dilution channel and total output, respectively.

The input concentrations of the primary odoants are *N* (*N* = 10, here) times of their corresponding optimal concentrations. The aim is to dilute the odorants to 1/*N* of the source concentration, when equally mixing *M* (*M* ≤ (*N*) odorants. It can be achieved by regulating the flow rate of the odorant channels to 1/*N* of the total flow, and the flow rate of dilution channel to 1 − *M*/*N* of the total flow. A simple solution is to just tune the flow rate of the dilution channel, while keep all other input and output channels flow constantly. With the flow rates measured from *F*1 and *F*2, we can feedback to regulate *P*2 to drive the flow rate *F*1 to its target value. In the process, *P*1 and *P*3 are kept as at preset values.

### Analysis of Neuronal Calcium Activity

#### Motion Correction

We collect volumetric recordings of multiple *Drosophila* larval neurons expressing GCaMP6m using a spinning disk confocal microscope. While the larva is well immobilized, motion relative to the size of a neuron persists during the recording and must be corrected. Depending on the extent of the motion and the neurons being imaged, we use a combination of rigid and piecewise rigid registration techniques to correct the motion. We typically perform an initial registration by spatially downsampling the recording and searching for translations that provide the best coarse alignment. If the neurons are densely packed, this registration may be sufficient. Otherwise, we may need to fine-tune the alignment either by rigid piecewise-rigid motion correction^37^.

#### Extracting calcium traces

We extract calcium traces from individual neurons within a multineuronal recording in different ways depending on the type of recording. For recordings of neuron cell bodies, we use the CaImAn package which performs constrained nonnegative matrix factorization^38,39^. This method performs well when the ROIs are similar in shape and spatially compact, but tends to perform worse on multineuron calcium recordings of neuropil, for example, in the larval antennal lobe. This is because ORN axons, PN dendrites, or other neurons that form synapses with a glomerulus in the AL form spatially distributed lobes.

To analyze such recordings (specifically for recordings of Keystone in this work), we developed a procedure for visualization of the temporal structure in a recording. We first remove background voxels using a manually set threshold. We then run t-SNE^19^ on the foreground voxels to embed the temporal structure into a low-dimensional space. This embedding relies only on the temporal information – spatial information is not used. We then cluster the points in the t-SNE embedded space using a density-based clustering algorithm^40^. In practice, the output from the clustering algorithm requires manual correction, and the t-SNE embedding provides easy visualization for that manual correction. Validation of the clustering is performed by observing the calcium time course for all voxels within a given cluster.

## Acknowledgements

The authors thank Jess Kanwal and Matthew Tu for testing the early prototypes of the microfluidic chip, Katrin Vogt for making the UAS-mCherry.NLS; UAS-GCaMP6m fly, and valuable discussions with Vivek Venkatachalam, Yu Hu, Luis Hernandez-Nunez and Cengiz Pehlevan. This work was performed in part at the Center for Nanoscale Systems (CNS), a member of the National Nanotechnology Coordinated Infrastructure Network (NNCI), which is supported by the National Science Foundation under NSF award no. 1541959. CNS is part of Harvard University. This work was also supported by an NSF Brain Initiative grant (NSF-IOS-1556388), and grants from the NIH (8DP1GM105383, P01GM103770). G.S. acknowledges support from the Harvard Brain Initiative Collaborative Seed Grant Program.

## Author contributions statement

G.S. conceived the project, designed the microfluidics and setup, conducted experiments, analyzed, interpreted the data and wrote the paper; J.B. designed the setup, developed the image analysis software, conducted experiments, analyzed, interpreted the data and wrote the paper; Y.F. designed the microfluidics; A.S. interpreted the data and wrote the paper.

---

**Figure S1.**
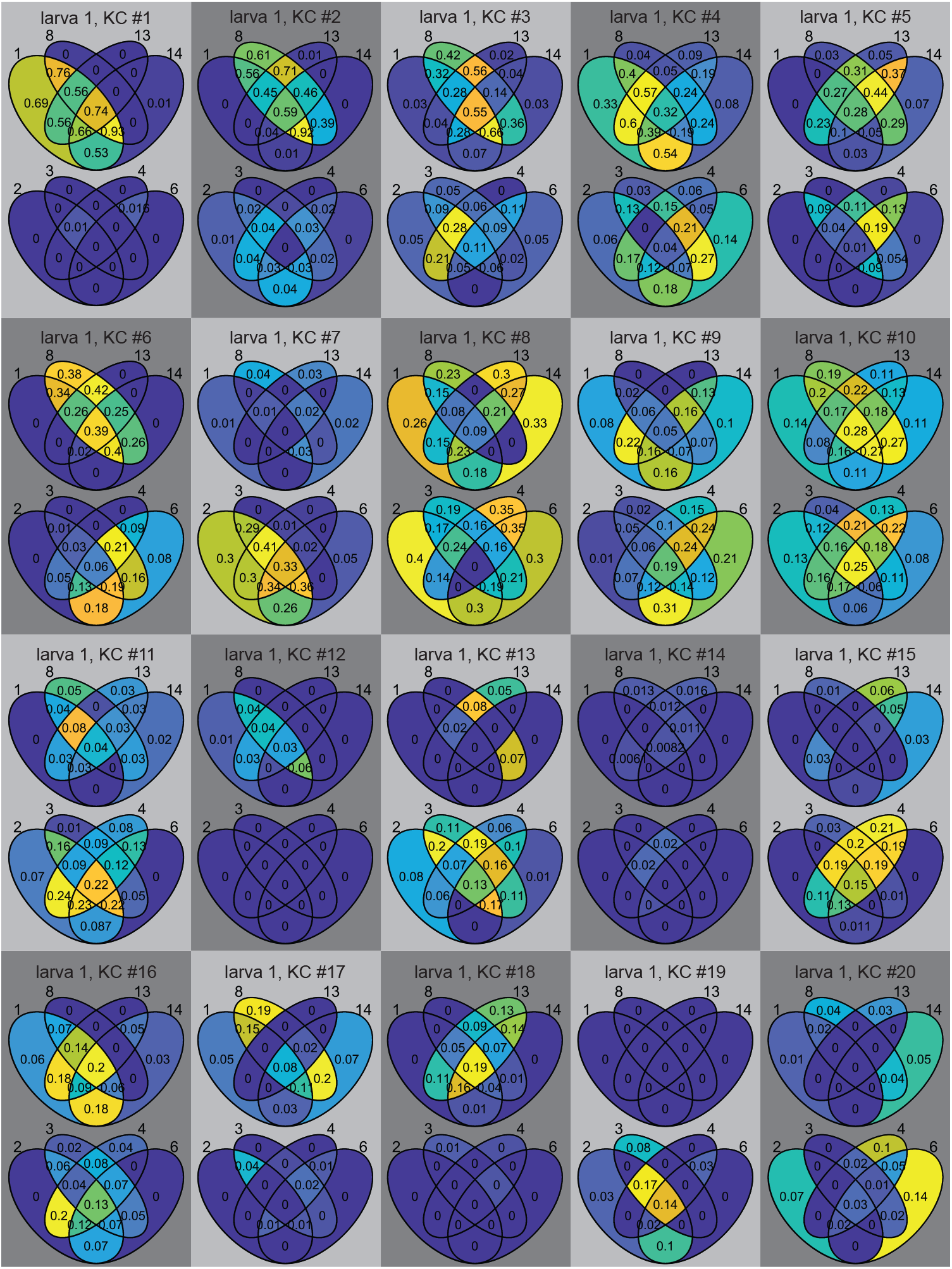
Visualizing ‘Animal 1’ KC responses to two complete set of primary odorant mixtures using Venn diagrams. Color bar for each KC’s plot is adjusted for its maximum response, to highlight the differences between the regions. The actual response value is labeled on the region.

**Figure S2.**
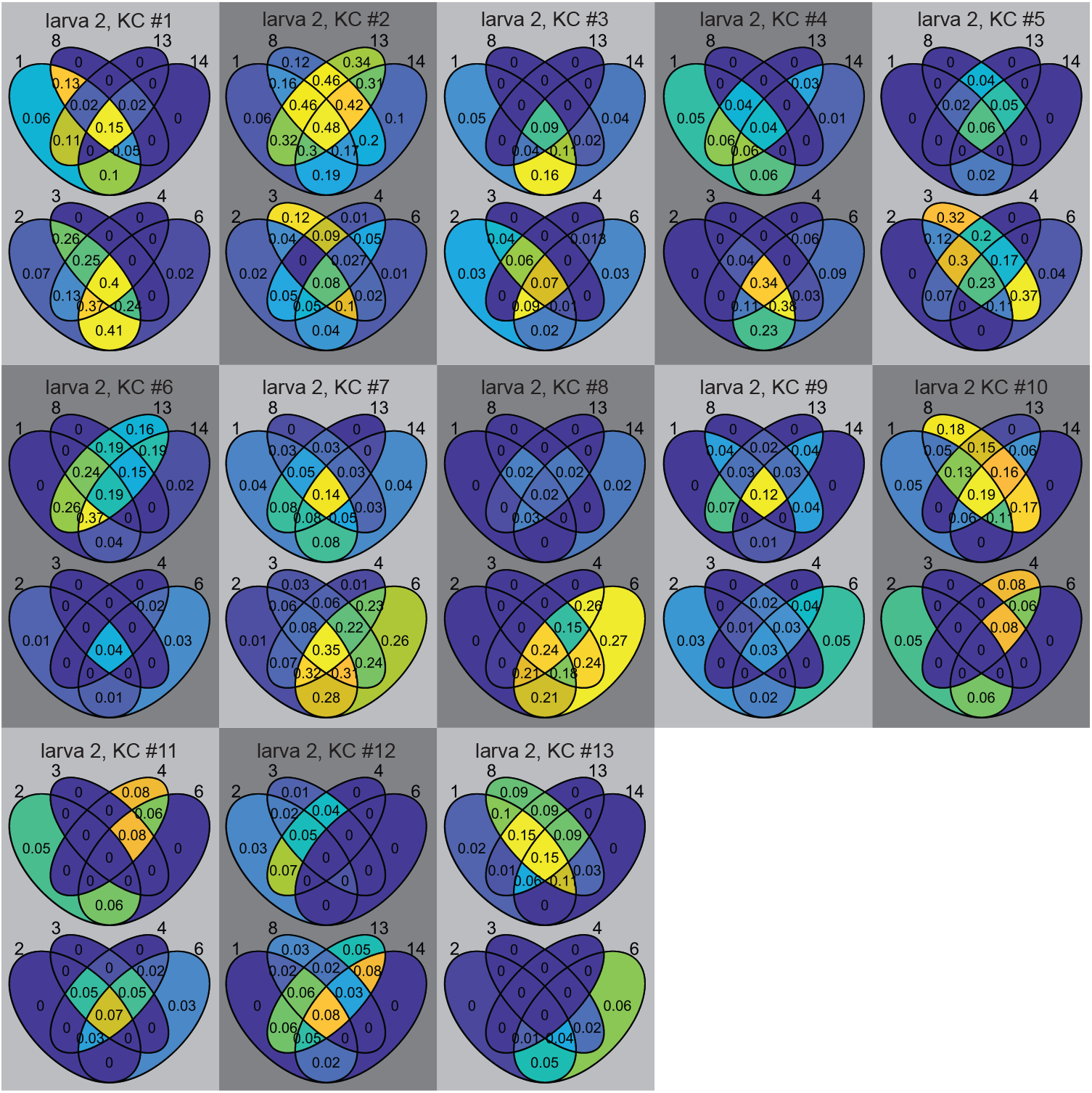
Visualizing ‘Animal 2’ KC responses to two complete set of primary odorant mixtures using Venn diagrams.

## References

1. Malnic, B., Hirono, J., Sato, T. & Buck, L. B. Combinatorial receptor codes for odors. Cell 96, 713–723 (1999).

2. Ramaekers, A. et al. Glomerular maps without cellular redundancy at successive levels of the *Drosophila* larval olfactory circuit. Curr. Biol. 15, 982–992 (2005).

3. Vosshall, L. B. & Stocker, R. F. Molecular Architecture of Smell and Taste in *Drosophila*. Annu. Rev. Neurosci. 30, 505–533 (2007).

4. Chronis, N., Zimmer, M. & Bargmann, C. I. Microfluidics for in vivo imaging of neuronal and behavioral activity in caenorhabditis elegans. Nat. Methods 4, 727 (2007).

5. Si, G. et al. Structured Odorant Response Patterns across a Complete Olfactory Receptor Neuron Population. Neuron 101, 950–962.e7 (2019).

6. Masse, N. Y., Turner, G. C. & Jefferis, G. S. Olfactory information processing in *Drosophila*. Curr. Biol. 19, R700–R713 (2009).

7. Hallem, E. A. & Carlson, J. R. Coding of Odors by a Receptor Repertoire. Cell 125, 143–160 (2006).

8. Kreher, S. A., Mathew, D., Kim, J. & Carlson, J. R. Translation of sensory input into behavioral output via an olfactory system. Neuron 59, 110–124 (2008).

9. Chen, T.-W. et al. Ultrasensitive fluorescent proteins for imaging neuronal activity. Nature 499, 295–300 (2013).

10. Vosshall, L. B., Amrein, H., Morozov, P. S., Rzhetsky, A. & Axel, R. A spatial map of olfactory receptor expression in the *Drosophila* antenna. Cell 96, 725–736 (1999).

11. Hong, C. C., Choi, J. W. & Ahn, C. H. A novel in-plane passive microfluidic mixer with modified Tesla structures. Lab on a Chip 4, 109–113 (2004).

12. Berck, M. E. et al. The wiring diagram of a glomerular olfactory system. eLife 5, e14859 (2016).

13. Bates, A. S. et al. Complete connectomic reconstruction of olfactory projection neurons in the fly brain. Curr. Biol. 30, 3183–3199 (2020).

14. Olsen, S. R., Bhandawat, V. & Wilson, R. I. Divisive normalization in olfactory population codes. Neuron 66, 287–299 (2010).

15. Olsen, S. R. & Wilson, R. I. Lateral presynaptic inhibition mediates gain control in an olfactory circuit. Nature 452, 956 (2008).

16. Chou, Y. H. et al. Diversity and wiring variability of olfactory local interneurons in the *Drosophila* antennal lobe. Nat. Neurosci. 13, 439–449 (2010).

17. Groschner, L. N. & Miesenböck, G. Mechanisms of Sensory Discrimination: Insights from *Drosophila* Olfaction. Annu. Rev. Biophys. 48, 209–229 (2019).

18. Li, H.-H. et al. A GAL4 driver resource for developmental and behavioral studies on the larval CNS of *Drosophila*. Cell Reports 8, 897–908 (2014).

19. Maaten, L. v. d. & Hinton, G. Visualizing data using t-SNE. J. Mach. Learn. Res. 9, 2579–2605 (2008).

20. Heisenberg, M. Mushroom body memoir: from maps to models. Nat. Rev. Neurosci. 4, 266 (2003).

21. Modi, M. N., Shuai, Y. & Turner, G. C. The *Drosophila* mushroom body: from architecture to algorithm in a learning circuit. Annu. Rev. Neurosci. 43, 465–484 (2020).

22. Murthy, M., Fiete, I. & Laurent, G. Testing Odor Response Stereotypy in the *Drosophila* Mushroom Body. Neuron 59, 1009–1023 (2008).

23. Caron, S. J., Ruta, V., Abbott, L. F. & Axel, R. Random convergence of olfactory inputs in the *Drosophila* mushroom body. Nature 497, 113–117 (2013).

24. Masuda-Nakagawa, L. M., Tanaka, N. K. & O’Kane, C. J. Stereotypic and random patterns of connectivity in the larval mushroom body calyx of *Drosophila*. Proc. Natl. Acad. Sci. 102, 19027–19032 (2005).

25. Eichler, K. et al. The complete connectome of a learning and memory centre in an insect brain. Nature 548, 175 (2017).

26. Gruntman, E. & Turner, G. C. Integration of the olfactory code across dendritic claws of single mushroom body neurons. Nat. Neurosci. 16, 1821–1829 (2013).

27. Li, H., Li, Y., Lei, Z., Wang, K. & Guo, A. Transformation of odor selectivity from projection neurons to single mushroom body neurons mapped with dual-color calcium imaging. Proc. Natl. Acad. Sci. 110, 12084–12089 (2013).

28. Pauls, D., Selcho, M., Gendre, N., Stocker, R. F. & Thum, A. S. *Drosophila* larvae establish appetitive olfactory memories via mushroom body neurons of embryonic origin. J. Neurosci. 30, 10655–10666 (2010).

29. Lin, A. C., Bygrave, A. M., De Calignon, A., Lee, T. & Miesenböck, G. Sparse, decorrelated odor coding in the mushroom body enhances learned odor discrimination. Nat. Neurosci. 17, 559–568 (2014).

30. Inada, K., Tsuchimoto, Y. & Kazama, H. Origins of Cell-Type-Specific Olfactory Processing in the *Drosophila* Mushroom Body Circuit. Neuron 95, 357–367.e4 (2017).

31. Jeanne, J. M., Fişek, M. & Wilson, R. I. The Organization of Projections from Olfactory Glomeruli onto Higher-Order Neurons. Neuron 98, 1198–1213.e6 (2018).

32. Chong, E. et al. Manipulating synthetic optogenetic odors reveals the coding logic of olfactory perception. Science 368 (2020).

33. Mathew, D. et al. Functional diversity among sensory receptors in a *Drosophila* olfactory circuit. Proc. Natl. Acad. Sci. 110, E2134–E2143 (2013).

34. Kuffler, S. W. Discharge patterns and functional organization of mammalian retina. J. Neurophysiol. 16, 37–68 (1953).

35. Burton, S. D., Wipfel, M., Guo, M., Eiting, T. P. & Wachowiak, M. A novel olfactometer for efficient and flexible odorant delivery. Chem. Senses 44, 173–188 (2019).

36. Gorur-Shandilya, S., Martelli, C., Demir, M. & Emonet, T. Controlling and measuring dynamic odorant stimuli in the laboratory. J. Exp. Biol. 222 (2019).

37. Pnevmatikakis, E. A. & Giovannucci, A. NoRMCorre: An online algorithm for piecewise rigid motion correction of calcium imaging data. J. Neurosci. Methods 291, 83–94 (2017).

38. Pnevmatikakis, E. A. et al. Simultaneous denoising, deconvolution, and demixing of calcium imaging data. Neuron 89, 285–299 (2016).

39. Giovannucci, A. et al. CaImAn an open source tool for scalable calcium imaging data analysis. Elife 8, e38173 (2019).

40. Biçici, E. & Yuret, D. Locally scaled density based clustering. In International Conference on Adaptive and Natural Computing Algorithms, 739–748 (Springer, 2007).

